# Palmitoylation controls the stability of 190 kDa Ankyrin-G in dendritic spines and is regulated by ZDHHC8 and lithium

**DOI:** 10.1101/620708

**Authors:** Nicolas H. Piguel, S. Sanders Shaun, Francesca I. DeSimone, Maria D. Martin-de-Saavedra, Emmarose McCoig, Leonardo E. Dionisio, Katharine R. Smith, Gareth M. Thomas, Peter Penzes

**Affiliations:** Department of Neuroscience, Northwestern University Feinberg School of Medicine, Chicago, IL 60611; Shriners Hospitals Pediatric Research Center, Lewis Katz School of Medicine at Temple University, Philadelphia, PA 19140; Instituto Universitario de Investigación en Neuroquímica, Department of Biochemistry and Molecular Biology, School of Pharmacy, Complutense University of Madrid, 28040 Madrid, Spain; Shriners Hospitals Pediatric Research Center and Department of Anatomy and Cell Biology, Lewis Katz School of Medicine at Temple University, Philadelphia, PA 19140; Department of Pharmacology, University of Colorado School of Medicine, Aurora, CO 80045; Department of Psychiatry and Behavioral Sciences, Northwestern University Feinberg School of Medicine, Chicago, IL 60611; Northwestern University Center for Autism and Neurodevelopment, Chicago, IL 60611

**Keywords:** ANK3, ankyrin-G, dendrite, dendritic spine, palmitoylation, lithium, ZDHHC8

## Abstract

AnkG, encoded by the *ANK3* gene, is a multifunctional scaffold protein with complex isoform expression: the 480 kDa and 270 kDa isoforms have roles at the axon initial segment and node of Ranvier, whereas the 190 kDa isoform (AnkG-190) has an emerging role in the dendritic shaft and spine heads. All isoforms of AnkG undergo palmitoylation, a post-translational modification regulating protein attachment to lipid membranes. However, palmitoylation of AnkG-190 has not been investigated in dendritic spines. The *ANK3* gene and altered expression of AnkG proteins are associated with a variety of neuropsychiatric and neurodevelopmental disorders including bipolar disorders and are implicated in the lithium response, a commonly used mood stabilizer for bipolar disorder patients, although the precise mechanisms involved are unknown. Here, we showed that Cys70 palmitoylation stabilizes the localization of AnkG-190 in spine heads and at dendritic plasma membrane nanodomains. Mutation of Cys70 impairs AnkG-190 function in dendritic spines and alters PSD-95 scaffolding. Interestingly, we find that lithium reduces AnkG-190 palmitoylation thereby increasing its mobility in dendritic spines. Finally, we demonstrate that the palmitoyl acyl transferase ZDHHC8, but not ZDHHC5, increases AnkG-190 stability in spine heads and is inhibited by lithium. Together, our data reveal that palmitoylation is critical for AnkG-190 localization and function and a potential ZDDHC8/AnkG-190 mechanism linking AnkG-190 mobility to the neuronal effects of lithium.

## Introduction

S-palmitoylation, a reversible form of post-translational modification regulating protein attachment to lipid membranes, is a critical process used to stabilize proteins in dendritic spines and dendrites (1) and localize proteins to different neuronal subcompartments (2). Important synaptic proteins, including neurotransmitter receptors, scaffold proteins, transporters, adhesion molecules, SNAREs, and other trafficking proteins, are palmitoylated (3) and palmitoylation also regulates dendrite and synapse morphology (4).

The *ANK3* gene encodes multiple isoforms of ankyrin-G (AnkG), of which the 190, 270, and 480 □kDa isoforms are the most prominent in the brain (5). The giant 270 and 480 kDa isoforms have been mostly studied for their function in the axon initial segment (AIS) (6) or nodes of Ravier (7). The 480 □kDa isoform has been shown to prevent gamma-aminobutyric acid (GABA) receptor endocytosis and to have an important role in maintaining GABAergic synapses (8). The 190 kDa isoform of AnkG is less well understood in neurons, but in contrast to larger isoforms, has been found to localize to dendritic spines and can be identified in the human postsynaptic density (PSD) fraction (9). Within subsynaptic nanodomains in dendritic spine heads and necks, AnkG-190 plays an important role in dendritic spine maintenance and long-term potentiation (10).

The membrane-binding regions of all three main AnkG isoforms harbor a cysteine residue (C70) that can be palmitoylated, a process that is required for AnkG association with the membrane, appropriate cellular localization, and function in non-neuronal cells (11, 12). Structural analysis has shown that the membrane anchoring of AnkG is facilitated by palmitoylation, defining a stable binding interface on the lipid membrane (13). Palmitoylation is known to play an important role in the function of 270 and 480 kDa isoforms, but the role of palmitoylation of the 190 kDa isoform in neurons is unknown (8, 11).

The human *ANK3* gene has been associated with various neuropsychiatric diseases, including bipolar disorder (BD), schizophrenia, and autism spectrum disorder (ASD) (14–16). Multiple independent genome-wide association studies (GWAS) have strongly linked *ANK3* to BD in a variety of different ethnicities (17–20) and have been corroborated by meta-analyses (21). In addition to a genetic association with *ANK3,* some studies have shown an increase in *ANK3* expression in the blood (22) and lymphoblastoid cells of BD patients (23), revealing the potential involvement of AnkG in disease mechanisms. Furthermore, a BD-related risk variant in *ANK3* is associated with decreased expression of a specific AnkG isoform in the cerebellum (24), and variants linked to loss of function are also implicated in ASD and intellectual disability (ID) (25). Thus, either increases or decreases in *ANK3* gene expression and isoform-specific variation may be implicated in the disorders. Lithium is a mood stabilizer commonly used to treat BD patients as a first-line intervention (26). Responsiveness of BD patients to lithium treatment appears to be a heritable trait (27) linked to genetic markers in patients isolated from GWAS (28). Interestingly, lithium has been used to correct BD-related behaviors and neuronal architecture deficits in Ank3 expression-deficient mouse models, suggesting a potential link between *ANK3* and the lithium response (29–32).

Understanding synaptic palmitoylation is likely to provide insights into a normal neuronal function but also pathophysiological and therapeutic mechanisms (33). Thus, we investigated the physiological function of AnkG palmitoylation and how it may be regulated by palmitoyl acyl transferases and a mood stabilizer.

## Materials and Methods

### Plasmids and antibodies

AnkyrinG-190-GFP was a gift from Vann Bennett (Addgene plasmid # 31059; http://n2t.net/addgene:31059; RRID:Addgene_31059). AnkG-mCherry was a gift from Benedicte Dargent (Addgeneplasmid #42566; http://n2t.net/addgene:42566; RRID: Addgene_42566). HA-ZDHHC5 and HA-ZDHHC8 were kind gifts from Dr. Shernaz Bamji. pEGFP-N2 and pmCherry-C1 (Clontech, Mountain View, CA, USA; now Takara Bio USA, Inc.). The following primary monoclonal and polyclonal antibodies (mAb and pAb, respectively) were used: AnkG mAb clone N106/20 (western blot) and clone N106/36 (immunocytochemistry) from Neuromab (Davis, CA, USA), PSD-95 (Neuromab), turboGFP mAb (to identify RNAi expression, Origene,), DsRed pAb (to identify mCherry expression, Clontech, Mountain View, CA, USA), and GFP chicken pAb (to identify GFP expression, Abcam, Cambridge, MA, USA).

### Neuronal cell culture and transfection

Dissociated cultures of primary cortical neurons were prepared from E18 Sprague-Dawley rat embryos. Brains were dissected in ice-cold Hank’s buffered salt solution, and cortical tissue was isolated, digested with papain (Sigma; diluted in Neurobasal with EDTA (0.5 mM) and DNaseI (2 units/mL), activated with L-cysteine (1 mM) at 37°C), and mechanically dissociated in neuronal feeding media (Neurobasal + B27 supplement (Invitrogen, Carlsbad, CA, USA) + 0.5 mM glutamine + penicillin/streptomycin). Dissociated neurons were plated at high density (300 000 cells/cm^2^) on 1.5 mm thickness polylysine-coated glass coverslips of 18 mm diameter (Warner Instrument, Hamden, CT, USA)). One hour after plating, the media was replaced. Neuronal cultures were maintained at 37°C in 5% CO_2_. The neuronal feeding medium was supplemented with 200 μM D,L-amino-phosphonovalerate (D,L-APV, Ascent Scientific, Bristol, UK) beginning on day 4 in vitro (DIV4). Neurons were transfected at DIV21 with Lipofectamine 2000 (Invitrogen), providing a transfection efficiency of 0.5-1%. Plasmids (2-5 μg total DNA). Briefly, Lipofectamine 2000 and DNA were diluted in Dulbecco’s Modified Eagle Medium + HEPES (10 mM), mixed thoroughly together, and incubated for 20-30 min at 37°C before addition to cultured cells in the absence of antibiotics. Following transfection, neurons were supplanted in antibiotic-containing feeding media containing half conditioned and half fresh media and allowed to express constructs for 3 days or as indicated.

### Immunocytochemistry

Cells were fixed for 10 min in 4% formaldehyde/4% sucrose in PBS and then in methanol pre-chilled to −20°C for 10 min. Fixed neurons were permeabilized and blocked simultaneously in PBS containing 2% normal goat serum and 0.2% Triton-X-100 for 1 h at room temperature. Primary antibodies were added in PBS containing 2% normal goat serum overnight at 4°C, followed by 3 x 10 min washes in PBS. Secondary antibodies were incubated for 1 h at room temp, also in 2% normal goat serum in PBS. Three further washes (15 min each) were performed before coverslips were mounted using ProLong antifade reagent (Life Technologies).

### Confocal microscopy

Confocal images of immunostained neurons were obtained with a Nikon (Amsterdam, Netherlands) C2 confocal microscope. Images of neurons were taken using the 63x oil-immersion objective (numerical aperture (NA) = 1.4) as z-series of 7-12 images taken at 0.4 μm intervals, averaged 2 times, with 1024×1024 pixel resolution. Detector gain and offset were adjusted in the channel of each cell fill (GFP or mCherry) to include all spines and enhance edge detection. Intensity plot profiles for dendrite/spine localization were performed in ImageJ. A 4 μm line was drawn across 3-5 spines per neuron and averaged across neurons to produce average intensity plot profiles ± SEM. Spine density, width, and length were analyzed with ImageJ. Epifluorescence images were obtained with a 10x objective, and traces of dendrites were drawn and analyzed with Sholl analysis in ImageJ. In culture, pyramidal neurons conserve their polarity and develop a long dendrite, which we refer to as the apical dendrite, and shorter dendrites near the soma, which we refer to as basal dendrites for Sholl analysis.

### Structured illumination microscopy imaging and analysis

Imaging and reconstruction parameters were empirically determined with the assistance of the expertise in the Nikon Imaging Center at Northwestern. The acquisition was set to 10 MHz, 14 bits with EM gain, and no binning. Auto-exposure was kept between 100-300 ms, and the EM gain multiplier was restrained below 300. Conversion gain was held at 1x unless necessary to increase the signal to 2.4x. Laser power was adjusted to keep max pixel intensity<4000 units. Three reconstruction parameters (illumination modulation contrast, high-resolution noise suppression, and out-of-focus blur suppression) were extensively tested to generate consistent images across experiments without abnormal features or artifacts and produce the best Fourier transforms. Reconstruction parameters (0.96, 1.19, and 0.17) were kept consistent across experiments and imaging sessions. 3D dendritic spine reconstruction was done on a NIS instrument (Nikon). Intensity plot profiling for dendrite analysis was performed with ImageJ (National Institutes of Health, *http://rsbweb.nih.gov/ii*). A punctate distribution at the membrane corresponds to a larger fluctuation in fluorescence intensity, which will result in a larger standard deviation (Puncta index).

### Pharmacological treatments

2-Bromopalmitate (2-BrP), a palmitoylation inhibitor, and S-methyl methanethiosulfonate (MMTS), a thiol blocker, were from Sigma (St. Louis, MO, USA). All other chemicals were from ThermoFisher Scientific (Waltham, MA, USA) and were of the highest reagent grade. 2-BrP was used at 20 mM, palmostatin B (which inhibits depalmitoylation) at 50 μM, lithium chloride at 2 mM, and all were applied for stimulation over 24 h.

### Acyl biotinyl exchange assay

For acyl biotinyl exchange (ABE) experiments, rat cortical neurons cultured as described above were lysed directly in buffer (50 mM HEPES pH 7.0, 2% SDS, 1 mM EDTA plus protease inhibitor mixture (PIC, Roche) and 20 mM MMTS (to block free thiols). Following lysis, excess MMTS was removed by acetone precipitation. Pellets were dissolved in a buffer containing 4% (wt/vol) SDS. Samples were diluted and incubated for 1 h in either 0.7 M hydroxylamine (NH_2_OH) pH 7.4 (to cleave thioester bonds) or 50 mM Tris pH 7.4, both containing sulfhydryl-reactive (HPDP-) biotin and incubated for 1 h at room temperature. Acetone precipitation was performed to remove unreacted HPDP-biotin and hydroxylamine and pellets were resuspended in lysis buffer without MMTS. SDS was diluted to 0.1% (wt/vol) and biotinylated proteins in the samples were affinity-purified using neutravidin-conjugated beads. Beta-mercaptoethanol [1% (vol/vol)] was used to cleave HPDP-biotin and release purified proteins from the beads. The released proteins in the supernatant were denatured in the SDS sample buffer and processed for SDS-PAGE. Adult rat forebrain was dissected, rapidly cooled in ice-cold recording buffer (11), and homogenized in 10 volumes of 4 mM HEPES, 320 mM sucrose, pH 7.4, containing fresh PIC and 20 mM MMTS. Samples were centrifuged to remove debris, brought to room temperature, and SDS was added to 1% (v/v) final concentration. Samples were then centrifuged at 27,000 x *g* for 30 min at 4°C and supernatants were subjected to acetone precipitation and ABE as described above.

### Fluorescence recovery after photobleaching

Neurons were imaged at 37°C, 5% CO_2_, with a Nikon C2 confocal microscope using a 63x oil immersion objective with NA = 1.4 in an OKOLAB stage-type CO_2_ incubator. Single plane images were captured with an EMCCD camera every 10 s for 200 s. 80% laser power pulses of 1 ms (2 iterations) were used to bleach GFP in the spine. After background subtraction and validation of the maximum 10% remaining fluorescence after photobleaching, data were normalized with the pre-bleach value. Recovery data points were then fitted to a one-phase association exponential in GraphPad Prism. The mobile fraction was calculated as an average of the plateaued fluorescence level and expressed as a percentage of the pre-bleached level.

### Statistical analysis

All statistical tests were performed with GraphPad Prism. Data were tested for normality with D’Agostino and Pearson methods to determine the use of non-parametric (Mann-Whitney, Kruskal-Wallis, Spearman correlations) or parametric (unpaired t-test, ANOVA, Pearson correlations) tests. Post-hoc tests were included in analyses with multiple comparisons. Bar graphs the mean ± SEM unless otherwise noted. Differences were considered significant if p≤0.05. N values refer to the number of cells per condition from a minimum of 3 biological replicates unless otherwise stated.

## Results

### Palmitoylation of C70 stabilizes AnkG-190 in dendritic spine heads and dendritic nanodomains

We have previously demonstrated AnkG-190 plays a key role in the dendritic spine and dendrite maintenance (10). However, the post-translational modifications that regulate these specific functions of AnkG-190 are unknown. Since palmitoylation of AnkG-190 in non-neuronal cells was shown to be essential for the association of AnkG with plasma membranes (11–13), we hypothesized that this modification may also modulate AnkG-190’s localization in dendrites. We first validated the presence of palmitoylated AnkG-190 in rat cortex utilizing the acyl-biotinyl exchange assay (11), which exchanges palmitoyl modifications to biotin (Fig. 1A). To prevent palmitoylation of AnkG-190, we mutated the cysteine 70 residue (11) to alanine (C70A) in a GFP fused construct (GFP-AnkG-190-C70A) and assessed how palmitoylation-deficient AnkG-190 could localize in neuronal dendrites and spines (Fig. 1B). Confocal images revealed that overexpressed GFP-AnkG-190 is enriched in the spine head, as previously observed (10). This enrichment is lost following C70A (Fig. 1C). Quantification revealed an abrogation in C70A spine head localization comparable to soluble GFP cell fill level (Fig. 1D). To access localization on AnkG-190-C70A in dendritic spines, we used super-resolution structured illumination microscopy (SIM) imaging (10, 34). Dendritic spine SIM imaging confirmed the results obtained by confocal microscopy. However, AnkG-190-C70A prevented the formation of these nanodomains present in the spine head observed when the AnkG-190 construct is expressed (Fig. 1E). SIM imaging of dendrites showed that the C70A point mutation did not alter the presence of AnkG-190 at the membrane but changed its localization pattern (Fig. 1F). AnkG-190-C70A was found to exhibit a more diffuse distribution pattern compared to AnkG-190 (Fig. 1F and G) measured by a smaller variation in fluorescence intensity (Puncta index). These results indicate that palmitoylation is important to maintain nanodomains within dendritic spines and along the dendrite.

**Figure 1.**
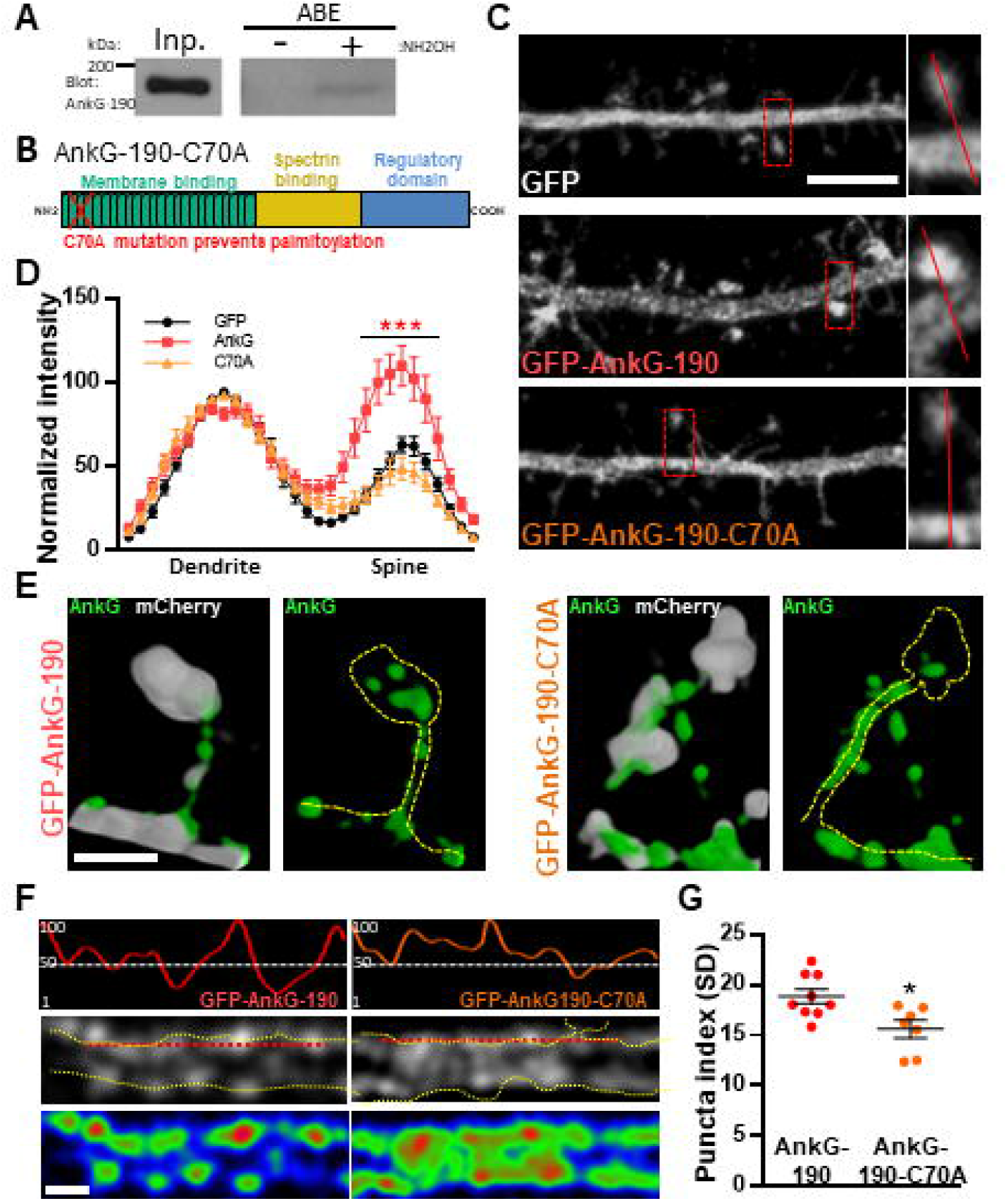
Palmitoylation of cysteine 70 stabilizes AnkG-190 in dendritic spines and dendrites. (**A**) AnkG-190 is palmitoylated in rat forebrain. Solubilized rat forebrain homogenates were subjected to ABE to purify palmitoylated proteins. ABE fractions and a portion of input homogenate (Inp.) were western blotted to detect AnkG. Exclusion of NH_2_OH was used as a control for assay specificity. (**B**) Schematic of 190 kDa Ankyrin-G isoform showing the mutated cysteine which prevents palmitoylation. (**C**) Confocal images of 24-day cultured cortical neurons expressing mCherry with GFP, GFP-AnkG-190, or GFP-AnkG-190-C70A for 1 day, scale=5 μm. (**D**) Linescan analysis of spine:dendrite ratio of expressed GFP, GFP-AnkG-190, or GFP-AnkG-190-C70A (16-20 neurons per condition, 2-way ANOVA with Dunnett’s post-test, ***p≤0.001, ±SEM). (**E**) Spine SIM imaging 3D reconstruction of cultured rat neurons expressing mCherry (white), GFP-AnkG-190, or GFP-AnkG-190-C70A (both green), scale=0.5 μm. (**F**) SIM imaging of dendrites from cultured cortical neurons expressing GFP-AnkG-190 or GFP-AnkG-190-C70A, scale=0.5 μm, in gray or pseudocolor (middle and bottom) and their associated normalized linescan (top). (**G**) Scatter plot of the puncta index average (7-9 neurons per condition, Mann Whitney, *p≤0.05, ±SEM) for GFP-AnkG-190 or GFP-AnkG-190-C70A overexpression.

### AnkG-190-C70A overexpression modifies dendritic spines density and PSD95 spine content

After establishing that the C70A mutation altered AnkG localization, we analyzed the impact of the palmitoylation-deficient AnkG-190 on dendritic spine morphology and PSD-95 content. We tested this by quantitative analysis of spine linear density and morphology in neurons coexpressing mCherry with wild-type or GFP-AnkG-190-C70A (Fig. 2A). Compared to GFP-AnkG-190, the mutated form had fewer dendritic spines which are associated with a longer neck (Fig.2B). We did not observe any effects of the mutation on spine width. AnkG-190 has been previously shown to increase the dendritic spine head (10) and we would expect an increase in synaptic scaffold proteins (35). Therefore, we decided to analyze PSD-95, a key component of post-synaptic density. Surprisingly, C70A mutation resulted in a robust reduction in PSD-95 accumulation in spine heads as compared to GFP-AnkG-190 (Fig. 2C and D). To assess whether the reduction of PSD-95 was associated with a change in spine size, we compare the size of PSD-95 puncta in the spine head with the area of the corresponding spine head. We observed a significant positive correlation between spine size and PSD-95 content (Fig 2E). Interestingly, this positive correlation was lost when neurons were transfected with AnkG-190-C70A suggesting the importance of palmitoylation for the AnkG-190 to maintain PSD-95 in the spine. Taken together, these data show that AnkG-190 palmitoylation is important to maintain dendritic spine density and PSD-95 in the spine head.

**Figure 2.**
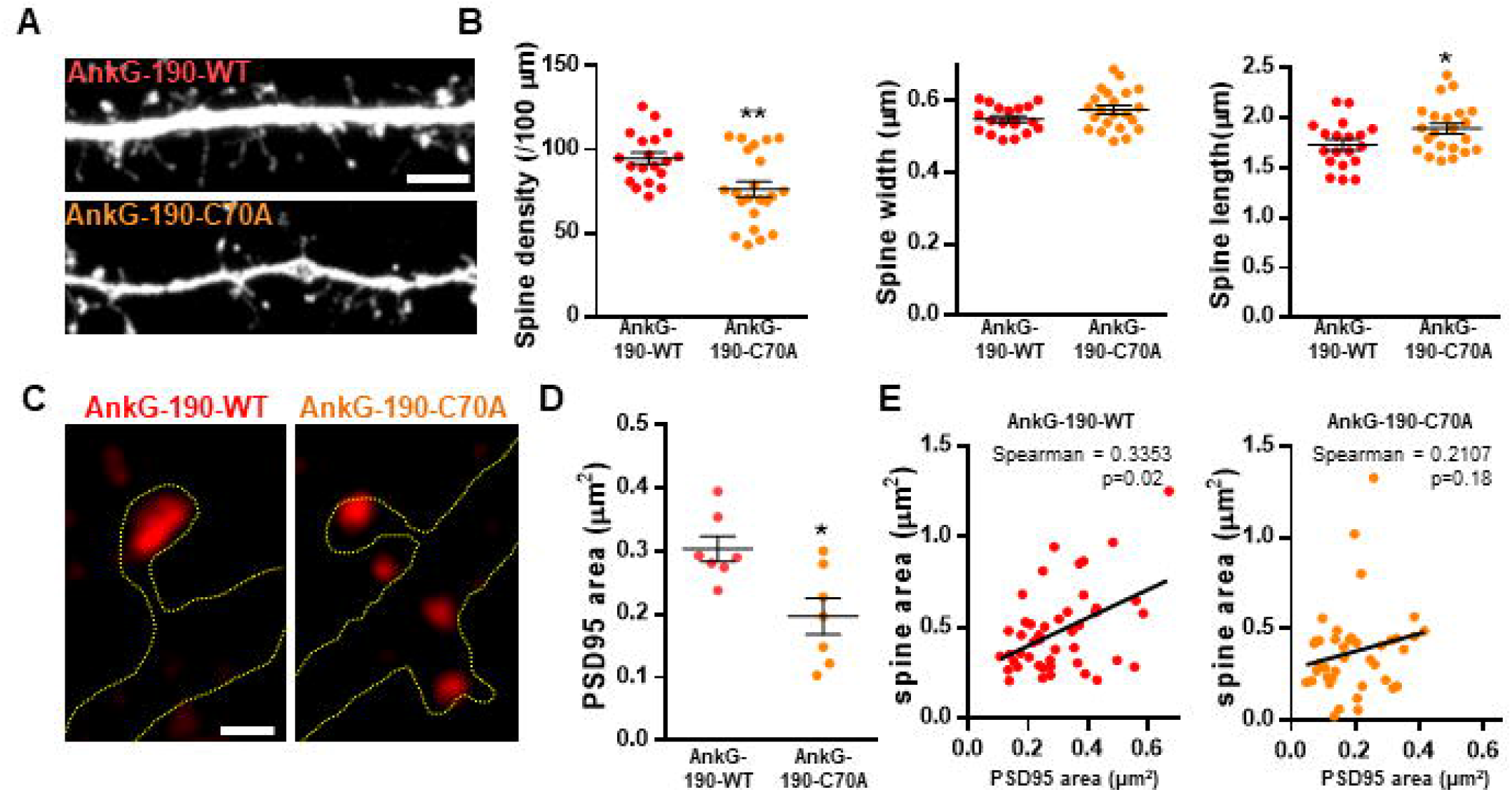
Effect of AnkG-190-C70A overexpression on dendritic spines. (A) Confocal images cultured cortical neurons expressing mCherry with GFP-AnkG190 or GFP-AnkG190-C70A, scale=5 μm. (B) Scatter plots showing quantification of spine density, spine width, and spine length for neurons expressing mCherry with GFP-Ank-G190 or GFP-AnkG-190-C70A (n=19-22 neurons, 1-way ANOVA with a Dunnett’s post-test, *p,0.05, *p,0.01, ***p,0. 001). (C) SIM images of PSD-95 from GFP-AnkG-190 and GFP-AnkG-190C-70A expressing, scale=0.5 μm. (D) Scatter plot graph showing a decrease of PSD-95 area in the spine with C70A mutation compared with WT (n=7 neurons, Mann Whitney, *p,0.05). (E) Correlation plot of spine area vs. PSD-95 area (40 to 50 spines) for GFP-AnkG-190 or GFP-AnkG-190-C70A overexpression.

### Lithium reduces AnkG-190 palmitoylation and increases its mobility

To investigate lithium-dependent modifications of AnkG, we performed an acyl biotinyl exchange (ABE) assay in cortical neuron cultures followed by western blotting of AnkG. Interestingly, treatment with lithium resulted in a ~58% decrease in palmitoylated AnkG-190 (Fig. 3A and B), a level similar to the negative condition containing the palmitoylation inhibitor 2-Bromopalmitate. On the other hand, lithium was not able to reduce GRIP1 palmitoylation (Fig S1A and B), another protein present in dendritic spines, demonstrating a level of specificity for AnkG-190. Consistent with a reduction in palmitoylation, immunocytochemistry of cortical neuron cultures after lithium treatment revealed a decrease in endogenous AnkG staining in mature spines (width ≥0.8 μm), which contain higher levels of AnkG (10), compared to vehicle treatment (Fig. 3C and D). These data demonstrate that lithium reduces the amount of AnkG localized in spines, which is consistent with the localization of palmitoylation-deficient AnkG. This result led us to hypothesize that lithium could prevent palmitoylation of AnkG-190 and therefore increase AnkG mobility in spines. To test this hypothesis, we performed fluorescence recovery after photobleaching (FRAP) in neurons overexpressing AnkG-190 and treated them with lithium for 24 h (Fig. 3E-G). Interestingly, lithium treatment increased the mobile fraction of AnkG-190 from ~52% to ~75% (Fig. 3G)comparable to the AnkG-190-C70A which cannot be palmitoylated. These data support an effect of lithium on the reduction of AnkG-190 palmitoylation leading to an increase in its mobility.

**Figure 3.**
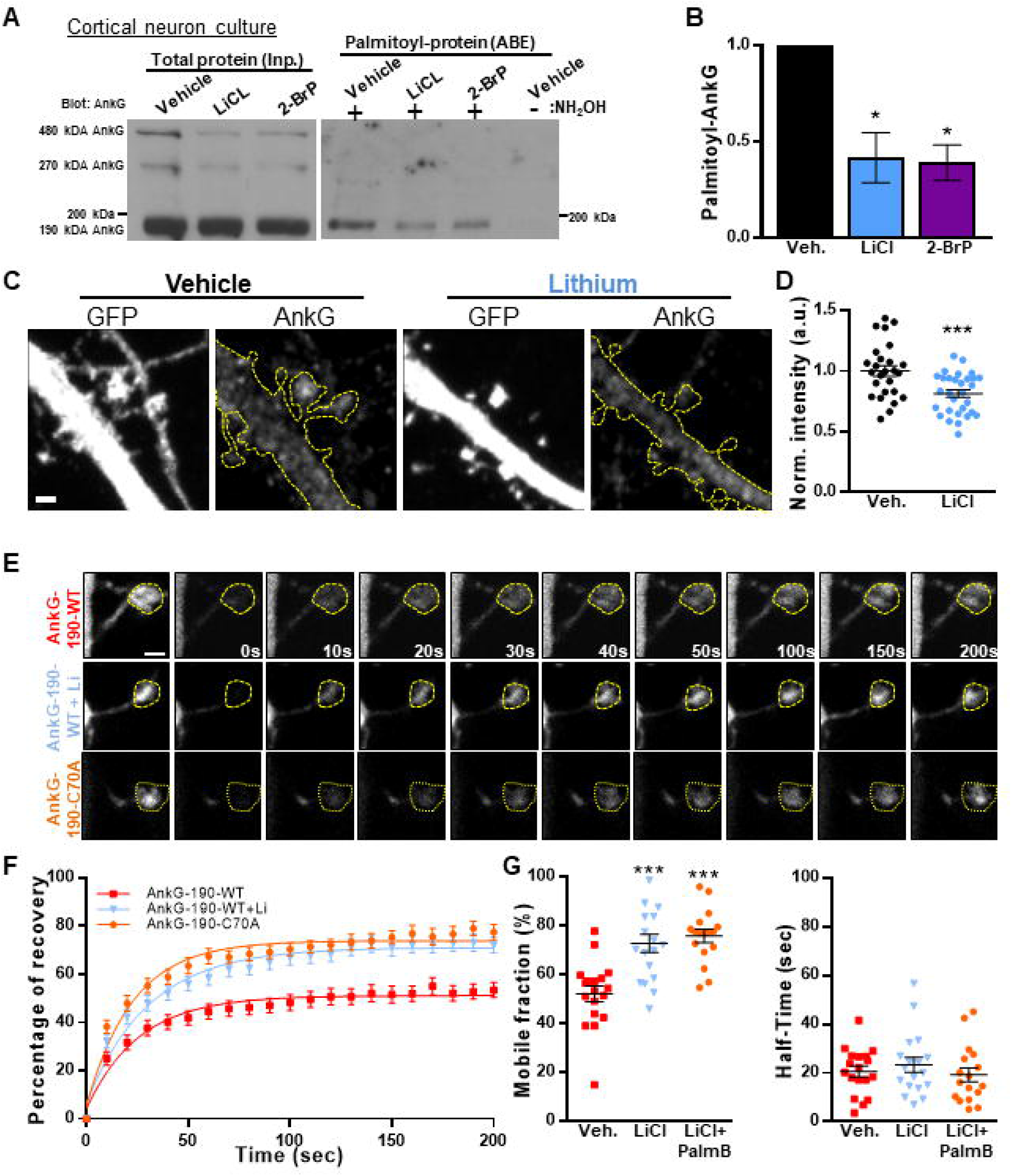
Lithium induces a decrease of palmitoylated AnkG-190 isoform and increases its mobility. (**A**) Lysates of cortical neurons treated with the indicated compounds were subjected to ABE to purify palmitoylated proteins. Levels of palmitoyl-AnkG-190 (top blot) and total AnkG expression in parent lysates (bottom blot) were detected with specific antibodies. Exclusion of NH_2_OH was used as a control for assay specificity. (**B**) Bar graph of 190 kDa AnkG isoform palmitoylation normalized with the input and relative to the untreated condition (5 independent experiments, t-test, *p≤0.05, ±SEM). (**C**) Representative maximal projection of a confocal image of endogenous AnkG staining in cortical neurons transfected with GFP with or without lithium stimulation, scale=2 μm. (**D**) Scatter plot of AnkG average intensity in mushroom spines on a single plane (26-30 neurons on 3 independent experiments, t-test, ***p≤0.001, ±SEM). (**E**) Representative time-lapse images of GFP-AnkG-190 fluorescence in 24 days rat neurons culture overexpressing GFP-AnkG-190 o GFP-AnkG-19-C70A for 3 days +/- lithium chloride during 1 day in FRAP experiments, scale=2 μm. (**F**) Quantification of GFP fluorescence in spines over time. Data are fitted with single exponentials (colored lines). Data are represented as mean ±SEM. (**G**) Scatter plots of mobile fraction (left) or half-time recovery (right) of GFP-AnkG-190 or GFP-AnkG-190-C70A +/- lithium chloride (n=15-22 neurons, 1-ANOVA with Dunnett’s post-test, ***p≤0.001)

### Lithium prevents ZDHHC8 to stabilize AnkG-190 in dendritic spines

Specificity in palmitoylation is achieved through the activity of various related palmitoyl acyl transferases (PATs) proteins. Lithium was found to decrease the palmitoylation of AnG-190, but not that of GRIP1, within neurons, suggesting a targeted effect of lithium on a subset of neuronal palmitoylated proteins. PATs that have the capability of palmitoylating AnkG have been described in HEK293 and MDCK cells and include ZDHHC5 and ZDHHC8. These PATs target the C70 residue on AnkG-190, but it is unknown whether a similar mechanism occurs in neurons (12). By using overexpression and FRAP imaging in neuron cultures, we assessed whether ZDHHC5 and/or ZDHHC8 were able to stabilize AnkG-190 in dendritic spines and if lithium could reverse the process. To ensure the spine mobility phenotypes were palmitoylation-dependent, we also treated cultures with Palmostatin B, an inhibitor of thioesterase APT1 and APT2 (36), which effectively block the depalmitoylation pathway. We found that ZDHHC5 did not affect the recovery of fluorescence of GFP-AnkG-190 (Fig. 4A-C) demonstrating that it does not modulate AnkG-190 mobility in spines. In contrast, ZDHHC8 significantly decreased the GFP-AnkG-190 mobile fraction in the dendritic spine (Fig. 4A, D, and E), suggesting it can palmitoylate AnkG-190 and stabilize the protein in the dendritic spine. Moreover, we found that lithium treatment can inhibit the effects of ZDHHC8 overexpression on GFP-AnkG-190 confirming its action on the palmitoylation process.

**Figure 4.**
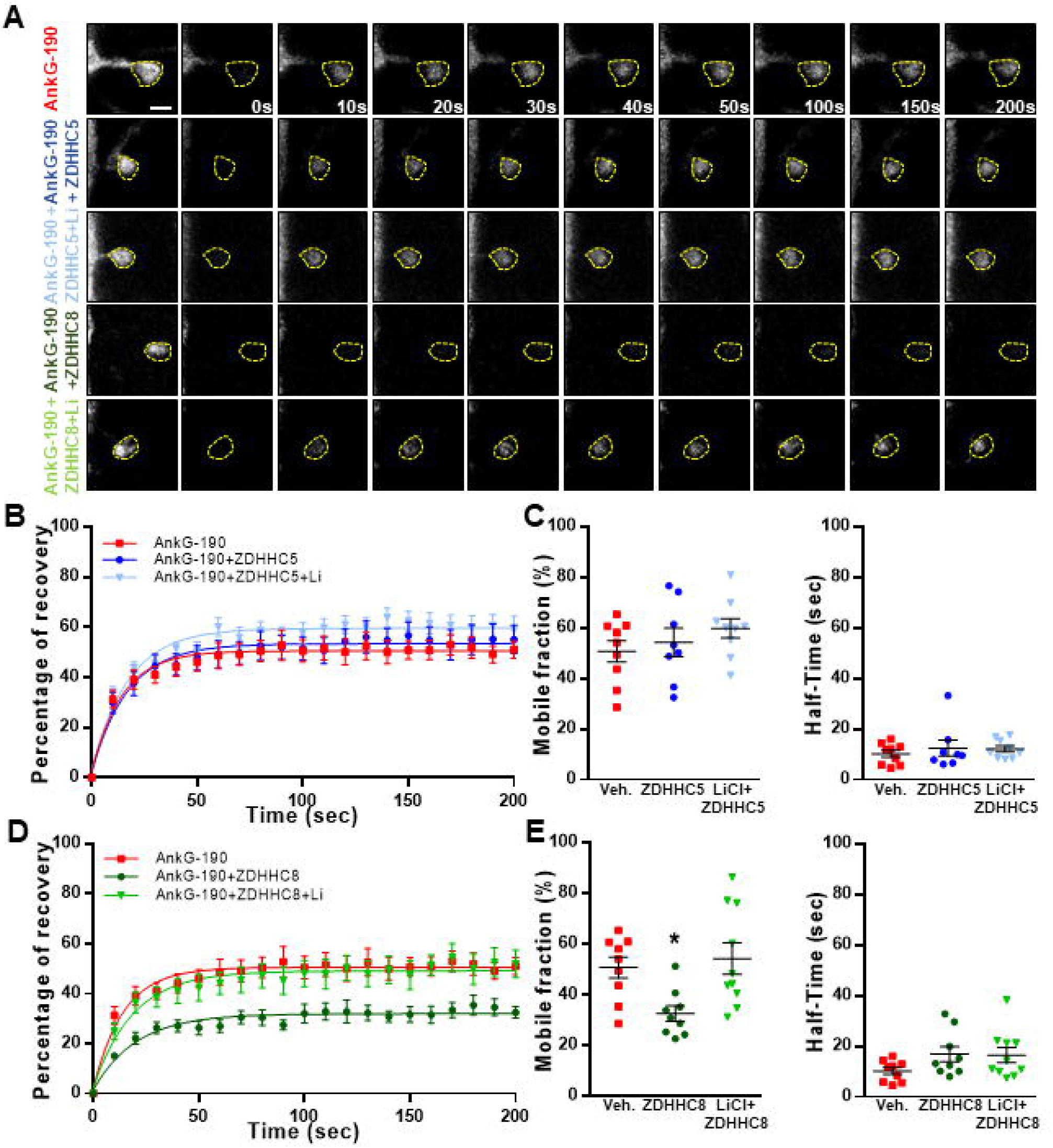
Lithium prevents DHHC8 to stabilize AnkG-190 in dendritic spines. (**A**) Representative time-lapse images of GFP-AnkG-190 fluorescence in 24 days rat neurons culture overexpressing GFP-AnkG-190 with or without DHHC5 or DHHC8 during 3 days in FRAP experiments, scale=2 μm. Cultures are treated with palmostatin B +/- lithium chloride 24 hours prior to the experiment (**B**) Quantification of GFP fluorescence in spines over time. Data are fitted with single exponentials (colored lines). Data are represented as mean ±SEM. (**C**) Scatter plots of mobile fraction (left) or half-time recovery (right) of GFP-AnkG-190 in the presence or absence of ZDHHC5 +/- lithium chloride (n=8-10 neurons, 1-ANOVA with Dunnett’s post-test). (**D**) Quantification of GFP fluorescence in spines over time. Data are fitted with single exponentials (colored lines). Data are represented as mean ±SEM. (**E**) Scatter plots of mobile fraction (left) or half-time recovery (right) of GFP-AnkG-190 in the presence or absence of ZDHHC8 +/- lithium chloride (n=8-10 neurons, 1-ANOVA with Dunnett’s post-test, *p≤0.05)

## Discussion

In the present study, we showed that palmitoylation on cysteine 70 stabilizes the localization of AnkG-190 in spine heads and at dendritic plasma membrane nanodomains. Remarkably, a mutant AnkG-190 lacking this palmitoylation leads to a reduction of PSD-95 in dendritic spine heads. We demonstrated that lithium treatment reduced AnkG-190 palmitoylation and increased its mobility in spines in a ZDHHC8-dependent pathway. Taken together, our data revealed a role for palmitoylation in dendritic spine components and uncovered a novel mechanism of lithium action on AnkG-190.

AnkG-190 shares most of the protein domain with the longer AnkG-480 isoform and should play similar roles in neurons. However, previous studies described a dendritic location of the AnkG-190 which can especially be found in dendritic spines (10) and PSD fractionation (9). Unlike the longer isoforms specific to the brain, AnkG-190 is expressed in other organs and has been studied in epithelial cells when it plays a role in lateral membranes (37, 38) and where it was associated with ßII-Spectrin (39, 40). Like in epithelial cells or for the neuronal AnkG-270 isoform (41), we showed that cysteine 70 is important for AnkG-190 localization in neuron dendrites. The C70A mutated AnkG-190 protein failed to form nanodomains in dendrites and dendrite spine heads. Our group showed previously that those nanodomains surround PSD-95 area (ref). Loss of AnkG-190 palmitoylation leads to a reduction of PSD-95 area without affecting the spine width suggesting AnkG-190 nanodomains stabilize PSD-95 but not the spine head architecture. Dendritic spine volume and PSD-95 size are linearly related (42) but the loss of PSD-95 is not a prerequisite for spine retraction (43) and AnkG-190 could serve as a link between the PSD and the cytoskeleton to maintain the correlation between the dendritic spine and PSD-95 volume. Although spine width does not significantly change, we can see a reduction in dendritic spine density and dendritic spine length, possibly due to a change of location AnkG-190 which will modify the localization of other proteins including dendritic spine initiators. We previously described a role for AnkG-190 in dendritic spine necks (10) and the absence of palmitoylation could push AnkG-190 neck location affecting the spine length.

We don’t lose AnkG-190 localization at the membrane or in the dendritic spine neck suggesting AnkG-190 can be stabilized by some protein partner interactions like spectrin (10). Unpalmitoylated AnkG has higher mobility within spines, indicating that palmitoylation restricts its mobility to specific sites, but still conserves a smaller stable population. These data suggest that only a subset of AnkG is palmitoylated at Cysteine 70 and other subpopulations of AnkG-190 exist in spines with distinct roles. Another possible partner that can maintain AnkG-190 in the dendritic spine is the scaffold in protein Homer1 that can bind Shank proteins to regulate spine morphology (44). Homer1 can bind directly PPXXF AnkG-190 motif through his EVH1 domain and mutation in the motif induces a loss of AnkG-190 localization and, like the mutation of the palmitoylation, does not affect all the immobile fraction (45).

Lithium has been shown to rescue some BD-related behavioral deficits in different *Ank3* KO mouse models (29–31), suggesting it can alleviate symptoms of mouse models with AnkG loss of function. However, heterozygous and homozygous loss of ANK3 likely represent rare mutations only present in a small subset of patients with ASD and ID, and the majority of individuals with BD are likely to retain AnkG expression or have increased expression (22, 23, 46). Moreover, the deletion of exon 37 targeting 270 kDa and 480 kDa isoforms induce a fivefold expression increase of the AnkG-190 isoform (47). Therefore AnkG-dependent mechanisms may also play a role in the lithium response. We show in neuron culture that lithium treatment has a drastic effect on AnkG-190 palmitoylation reducing it at the same level as the 2-BrP condition, suggesting lithium blocks all the unpalmitoylated AnkG-190 to be palmitoylated. Furthermore, the reduction of endogenous AnkG in the spine head and increase of GFP-AnkG-190 mobile fraction in the dendritic spine supports the reduction of palmitoylation observed.

Only ZDHHC8, one of the two PATs described to palmitoylate AnkG-190 (11), can stabilize GFP-AnkG-190 in the dendritic spine when we overexpressed it. This effect was abolished after 24 hours of lithium stimulation supporting the decrease of AnkG-190 palmitoylation due to the inhibition of ZDHHC8 activity. Moreover, palmitoylated GRIP1, which is a target of ZDDHC8 too (11), is not decreased during lithium treatment indicating a specific inhibition of AnkG-190 palmitoylation. We cannot exclude that ZDHHC5 affect AnkG-190 in other neuron subdomains to alter different function like the specific effect of ZDHHC8 on PICK1-dependent LTD (33). While *ZDHHC8* was not yet associated with BD (48), it has been related to schizophrenia (ref) and epilepsy (49) overlapping with ANK3 (50).

Our work describes a new mechanism of mood stabilizer action and its effects on two important psychiatric disorder risk factors *ANK3* and *ZDHHC8.* These findings open new directions for understanding basic control of neuroarchitecture but may also provide new opportunities for novel therapeutic targets.

## Footnotes

*To whom correspondence should be addressed.

## Author contributions

N.H.P., G.M.T., and P.P. designed research; N.H.P., F.I.D., S.S.S., R.G., K.E.H., L.E.D., and K.R.S. performed research; N.H.P., F.I.D., S.S.S. analyzed data; N.H.P and P.P. wrote the paper.

The authors declare no competing interests.

## Acknowledgments

This work was supported by R01MH107182 to P.P. Imaging work was performed at the Northwestern University Center for Advanced Microscopy, generously supported by NCI CCSG P30 CA060553 awarded to the Robert H. Lurie Comprehensive Cancer Center. We are grateful to members of the Penzes lab for helpful discussions, especially Dr. Marc P. Forrest, Dr. Marc Dos Santos, Dr. Seyoun Yoon, and Dr. Euan Parnell for his help on the manuscript. Dr. Antonio Sanz-Clemente is gratefully acknowledged for providing access to his lab equipment. All experiments involving animals were performed according to the Institutional Animal Care and Use Committee of NU.

## Notes

### Competing Interest Statement

The authors have declared no competing interest.

### Summary of Updates

Old Figure 1 removed Old Figure 2 revised New Figure 2 Author updated The manuscript text was updated to reflect the changes.

