## Supplemental Figure 1 for "Palmitoylation controls the stability of 190 kDa Ankyrin-G in dendritic spines and is regulated by ZDHHC8 and lithium"

### Figure Supp. 1

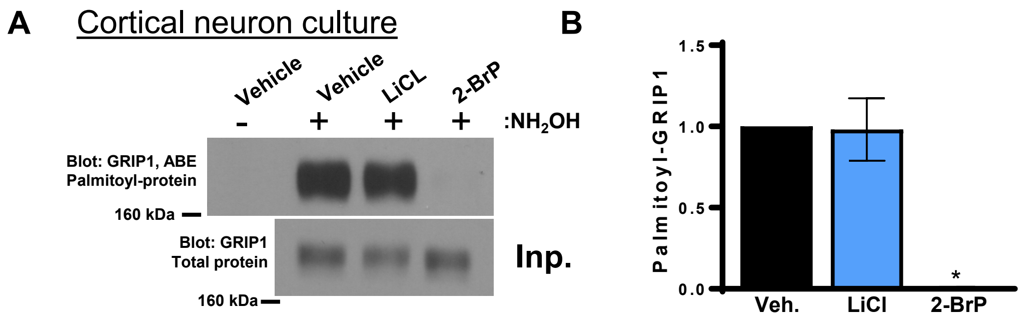


**Figure S3.** (**A**) Lysates of cortical neurons treated with the indicated compounds were subjected to ABE to purify palmitoylated proteins. Levels of palmitoyl-GRIP1 (top blot) and total AnkG expression in parent lysates (bottom blot) were detected with specific antibodies. Exclusion of NH2OH was used as a control for assay specificity. (**B**) Bar graph of GRIP1 palmitoylation normalized with the input and relative to the untreated condition (4 independent experiments, Kruskal-Wallis with Dunn's post-test, *$p \leq 0.05$, ±SEM).
